# The phonology of sperm whale coda vowels

**DOI:** 10.1101/2025.06.09.658556

**Authors:** Gašper Beguš, Maksymilian Dabkowski, Ronald L. Sprouse, David F. Gruber, Shane Gero

## Abstract

In previous research, sperm whale *codas* (structured series of clicks used for communication) have been shown to resemble human vowels acoustically. Based on the number of formants, two different coda quality categories have been described: *a*-codas and *i*-codas. In the present paper, we demonstrate that sperm whale codas not only resemble human vowels acoustically, but also pattern like them across several dimensions. First, traditional count- and timing-based coda types interact with coda “vowel” quality (*a* vs. *i*). Second, *a*-codas are generally longer than *i*-codas. Third, the duration of *i*-codas has a bimodal distribution, showing a contrast between short *i*-codas and long *ī*-codas. Fourth, the baseline coda length differs across whales. And fifth, edge clicks mismatching their coda often match an adjacent coda, a phenomenon that resembles human coarticulation. All five properties have close parallels in the phonetics and phonology of human languages. Sperm whale coda vocalizations thus represent one of the closest parallels to human phonology of any known animal communication system.

**SIGNIFICANCE STATEMENT:** Sperm whales communicate using series of clicks known as *codas*. The codas acoustically resemble human vowels. In addition, they pattern in ways similar to human sound systems. For example, different coda types are correlated with particular click qualities, and their durations are intentionally controlled. This shows that sperm whale vocalizations are highly complex and likely constitute one of the most sophisticated communication systems in the animal kingdom. By studying it, we may be able to gain a broader understanding of animal intelligence and social behaviors, determine the impact of human activities on whale habitat, develop strategies to protect whales from threats such as noise pollution and ship traffic, and advance legislation which facilitates that protection.

## 1 Introduction

Sperm whales (*Physeter macrocephalus*) are large pelagic mammals that live in small, matrilineal social units that belong to cultural clans characterized by social complexity and sophisticated behavior.^1^ To communicate, sperm whales use short series of clicks known as *codas*, which play a crucial role in maintaining social bonds and coordinating activities.^1–5^In addition, different clans of whales use different coda types,^2,6–11^suggesting that at least some aspects of the vocalization system are acquired (not innate).^2,12^

Traditionally, the sperm whale codas have been analyzed as grouped into different *coda types* based on the number of clicks and the length of inter-click intervals (ICI).^13^ Recent work reveals previously unobserved complexities in the timing features of the sperm whale communication system. Sharma *et al*. (2023) show that the traditional coda types are formed from four semi-independent parameters: rhythm (the relative timing between clicks in a coda), tempo (the duration of the coda), ornamentation (the addition of an extra click to a coda), and rubato (the modulation of tempo across consecutive codas in an exchange).^14^

Beguš *et al*. (2024) have recently opened a new dimension in the study of sperm whale clicks: their spectral properties. In particular, Beguš *et al*. show that whale clicks have structured spectral properties that (i) fall into one of two discrete categories (one vs. two formants), (ii) are orthogonal to the traditional coda type distinctions, (iii) recur across individual sperm whales, and (iv) are most likely actively controlled by the whales. They refer to the one-formant codas as “*a*-coda vowels” and to the two-formant codas as “*i*-coda vowels.”^15^

Despite a growing number of reports of the sperm whale communication system’s structural complexity, and potential expressivity, little is known about what information is conveyed by sperm whale codas, nor how exactly it may be encoded within their click-based system. Consequently, sperm whale vocalizations remain one of the most intriguing communication systems in the animal kingdom, and their structure presents an open question in animal research.^6,15^

In this paper, we explore the structural properties of whale codas and draw parallels to phenomena in human phonology. We argue that coda vowels are not mere acoustic resemblances of human vowels, but also behave similarly to human vowels across several dimensions.^1^ Specifically, we show that (i) there is a correlation between ICI-based coda *type* and coda vowel category a. k. a. *quality* (i. e. *a*-vs. *i*-vowels), (ii) *a*-vowels are generally longer than *i*-vowels, (iii) *i*-vowels have a bimodal distribution in their duration, (iv) different whales have different baseline coda durations, and (v) edge clicks mismatching the quality of their coda often match the quality of an adjacent coda. We find that all five features of our dataset have close structural analogues to phonetic and phonological patterns found in human languages. Thus, sperm whale vocalizations represent—to the best of our knowledge—one of the closest parallels to human phonology of any animal communication system.

## 2 Data

Our dataset consists of coda vocalizations produced by social units of Eastern Caribbean sperm whales encountered largely off the coast of the Island of Dominica (EC1). All of the recordings were collected from 2014 to 2018 by The Dominica Sperm Whale Project.^16^ The data includes recordings from a total of fifteen female and immature sperm whales. (Mature male sperm whales live predominantly solitary lives.^17^) In total, the data set contains 3948 temporally-ordered, speaker-identified, and manually annotated codas. Thus, the analyses we present are based on one of the largest datasets of sperm whale vocalizations produced by tagged whales. For more details on the dataset, including a discussion of the coda annotations, see Beguš *et al*. (2024).^15^

All audio has been captured with DTAG-3s—digital acoustic recording tags which are minimally invasive and use built-in hydrophones to record audio continuously at sampling rates of either 120 or 125 kHz with a 16-bit resolution.^18^ The analyzed vocalizations come from single-channel recordings (either left-channel or right-channel). All the codas were classified as either *focal*, i. e. produced by the whale carrying the DTAG that captured the sound, or *non-focal*, i. e. produced by a whale different from that carrying the DTAG. In all our analyses, only the focal codas have been used. This is to say, codas produced by other (non-focal) whales during exchanges were excluded. This ensured that the distance between the hydrophone and the sound source was the same in all the recordings within the same tag deployment, which allowed us to control for the acoustic distortions potentially created by the underwater environment. For a further discussion of the controls, see Beguš *et al*. (2024), who described and analyzed the same dataset of EC1 clan recordings.^15^

We have labeled the codas by hand based on the number of observed formats. Codas with one formant were labeled as “*a*” and codas with two formant as “*i*.” Codas that were not recorded clearly enough to allow for hand-labeling were excluded. After the exclusion of non-focal and noisy tokens (where by *tokens* we mean individual coda productions), we were left with 1144 codas. In some of the analyses, we applied further exclusion criteria, which are detailed in subsections 4.1-4.5.

## 3 Background

Sperm whale codas have traditionally been classified based on the number of clicks and the length of inter-click intervals.^13^ Different coda types have been given labels such as 3R, 3D, 4R_1_, 4R_2_, 5R_1_, 5R_2_, 5R_3_, 7D, 8D, 8i, 9i, 1+1+3, etc. The digits stand for the number of clicks, “R,” “D,” and “i” stand for “classes” of codas based on changes to the inter-click interval duration, which is—respectively—*regular, decreasing*, or *increasing* over the course of the coda, and the plus symbol “+” indicates an additional pause after a click. (Subscripts differentiate variants of the same coda rhythm, but with increasing duration.^9^) As such, for example, “1+1+3” stands for a coda that consists of two clicks followed by pauses and three clicks in quick succession (click…click…click-click-click) (illustrated in Figure 1a,d),^2^ “5R” stands for a coda that consists of five evenly spaced clicks (click-click-click-click-click) (Figure 1b,e) (with 5R_1_ codas being considerably are shorter than 5R_2_ codas), and “9i” stands for a coda that consists of a nine-click with increasing ICIs between clicks (Figure 1c).^9, 3^

**Figure 1.**
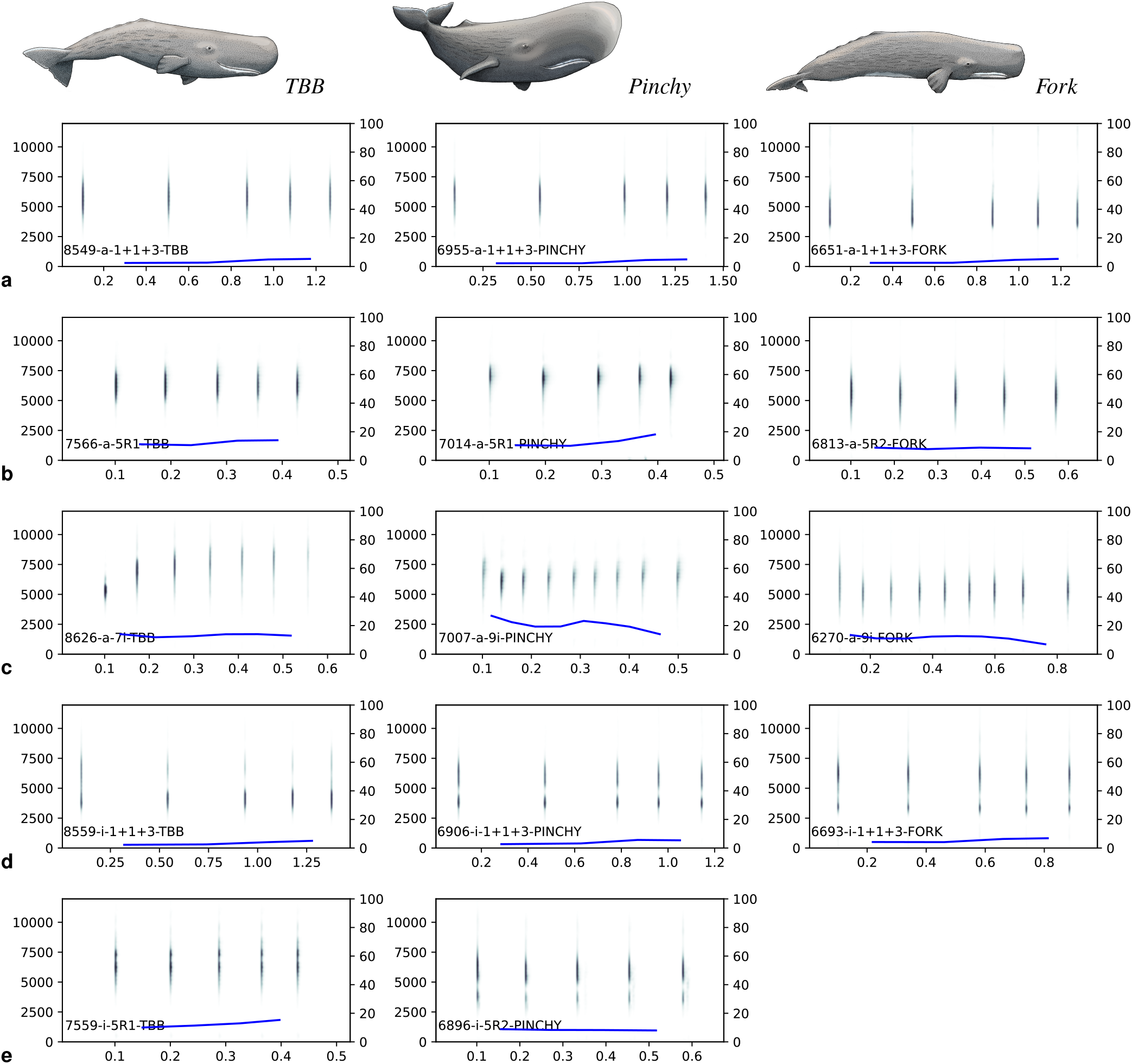
Different coda vowel quality and type combinations as produced by three different whales, including *a*-quality codas of the 1+1+3 type (**a**), 5R type (**b**), and i-class (“increasing” ICI) types (**c**), as well as *i*-quality codas of the 1+1+3 type (**d**) and 5R type (**e**). Pitch plots are given as blue lines.

Human vowels consist of a sequence of glottal pulses produced by vocal folds. Whale codas consist of a sequence of clicks produced by vibrating phonic lips, which play a role similar to the human vocal folds.^15^ In human languages, the frequency of glottal pulses corresponds to pitch—closely spaced glottal pulses give rise to a higher pitch, while more widely spaced pulses give rise to a lower pitch. In linguistics, *tone* refers to pitch as recruited to express linguistic meaning. Many languages use tone to distinguish between different words. For example, in Mandarin Chinese, the following four words differ only in their tonal contour, while having the same consonants and vowels:^25^ high and level tone *mā* ‘mother,’ rising tone *má* ‘hemp,’ falling-rising tone *ma?* ‘horse,’ and falling tone *mà* ‘scold.’ The coda types are therefore analogous to human tone: “regular” coda types can be compared to level tones, codas with “increasing” inter-click intervals are falling tones, and codas with “decreasing” inter-click intervals are rising tones.^4^ In Figure 1, the “pitch” of each coda is represented with a blue line.^5^

Additionally, the sperm whale communication system appears to distinguish between codas with the same rhythm type but different number of clicks, e. g. 7D vs. 8D or 8i vs. 9i. This has an analogy in systems where different tonal contours have inherent length, such as the Vietnamese contrast between short (*sac, nang*) and long (*nga, hoi*) tones.^26^

Beguš *et al*. (2024) additionally observe that the clicks which make up sperm whale codas have discrete spectral properties— each click has either one or two formants. Building on this observation, they propose to apply the source-filter model^27^ (originally formulated to explain the articulatory mechanics and acoustic properties of human speech) to whale coda clicks as well. Specifically, they speculate that clicks are produced by an interaction of vibrating phonic lips (the sound source, equivalent to human vocal folds) and the distal air sac (the acoustic filter, equivalent to the human vocal tract). Sperm whale coda clicks are thus analogized to human vocal pulses.

The two discovered categories of clicks differ in the number of formants: some clicks have one, while others have two. In human speech, the frequency of broad peaks in the spectrum (referred to as *formants*) determines which vowel is perceived. Given the similarities in the mechanisms of production and the acoustic properties between the human voice and whale vocalizations, Beguš *et al*. (2024) call them *coda vowels*. Codas with one-peak clicks are referred to as *a*-vowels (illustrated in Figure 1a-c), while those with two-peak clicks are referred to as *i*-vowels (Figure 1d-e). Here, we will refer to the contrast between *a*- and *i*-vowels as a difference in vowel *quality*. Furthermore, some of *a*-vowels show a dynamic formant trajectory, with their formant rising or lowering over the course of the coda (Figure 1c). Beguš *et al*. (2024) analogize these to human language diphthongs, such as in the English word *site* /*sajt*/, which likewise show vowel-internal formant transitions.^15^

Vowel quality is most likely actively controlled by whales. First, vowel quality has a clear bimodal distribution, with clicks showing either one or two formants. Second, the two types are discretely distributed across codas, i. e. there is little mixing of different click qualities within a coda (i. e. they are typically either all *a*-clicks or all *i*-clicks). Third, the rising or falling trajectories of resonant frequencies in “diphthongs” (Figure 1c) are similar across different coda tokens. Fourth and last, the frequencies do not appear to be crucially affected by movement, and the described patterns are observed across different individual whales.^15^ Beguš *et al*. (2024) show that different coda vowel qualities can be instantiated on the same coda types, and propose that coda type and coda quality are orthogonal.^15^ This points to another parallelism between the sperm whale communication system and human language, as tone and vowel quality are often similarly orthogonal. For example, in Mandarin Chinese, the falling-rising tone may appear on any vowel, e. g. *ma?* ‘horse,’ *m?i* ‘rice,’ *mo?* ‘smear,’ etc. Our paper builds on Beguš *et al*.’s

(2024) findings and reveals further complexities within the system of sperm whale vocalizations.

## 4 Results

### 4.1 Vowel quality–coda type correlation

First, we demonstrate that although the same coda type may have different vowel qualities, coda vowels show up unevenly across coda types. Throughout the paper, codas are categorized as “*a*” or “*i*” based on hand-labeling.^6^ Additionally, we exclude all coda types that had fewer than 15 tokens in the database. (In statistical models where codas with fewer tokens per type are included, the coefficients are difficult to estimate.) After the exclusion, we were left with 1066 codas, which were used in this analysis. We observe that in the 1+1+3 type, approximately half of the codas are of the *a*-coda type, and half are of the *i*-coda type.

The only other coda type with an approximately even distribution of coda vowels is 6-UNCLASSIFIED (which is a collection of different 6-click codas that could not be clustered into a defined type). In other coda types, such as 5R_1_ or 5R_2_, the *i*-coda vowel is attested but rarer than the *a*-coda vowel. In some coda types, the *i*-coda vowel is entirely unattested, such as the 8i, but given the low number of tokens of these coda types, it is possible that this is the result of undersampling.

To confirm these observations, we have fit the data to a mixed-effect logistic regression model with manually annotated coda vowels (*a* vs. *i*) as the dependent variable and coda type as the independent variable with a random intercept for each whale identity. While the distribution of *a*- and *i*-coda vowels is not significantly different from 50% in the 1+1+3 (*β* = − 0.20, *z* = − 1.04, *p* = 0.3) and the 6-UNCLASSIFIED coda types, the *i*-coda vowel is significantly less frequent on 5R_1_, 5R_2_, 6i, and 9i coda types. All models’ coefficients are given in Table 1.^7^ The distribution of the coda vowels is summarized in Figure 2.

**Table 1.** Coefficients of the mixed effects logistic regression model with the manually annotated *a*-vs. *i*-coda vowel as the dependent variable (*a* is coded as 1) and coda type as the dependent variable (treatment-coded with 1+1+3 as the base level). The model includes a random intercept for whale identity.

|  | estimate | std. error | z-value | Pr(>\|z\|) |
| --- | --- | --- | --- | --- |
| (intercept) | -0.20 | 0.19 | -1.04 | 0.2963 |
| coda type 5R<sub>1</sub> | -2.33 | 0.28 | -8.17 | 0.0000 |
| coda type 5R<sub>2</sub> | -2.22 | 0.61 | -3.62 | 0.0003 |
| coda type 6-UNCLASS. | 0.77 | 0.39 | 2.00 | 0.0457 |
| coda type 6i | -3.07 | 1.03 | -2.97 | 0.0030 |
| coda type 8i | -22.77 | 114.49 | -0.20 | 0.8424 |
| coda type 9i | -2.32 | 0.75 | -3.10 | 0.0020 |

**Figure 2.**
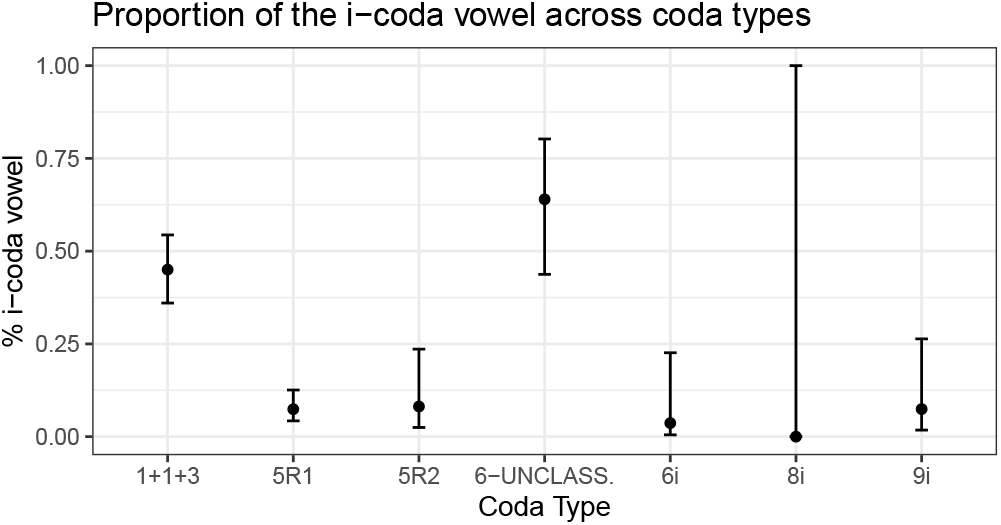
Proportion of the *i*-coda vowel across coda types. Estimates are from the mixed effects logistic regression model random intercept for whale identity. in Table 1 with 95% CIs.

Thus, we see that the two coda vowel qualities are evenly distributed in some coda types, while in other coda types, the *a*-vowel is much more frequent. Beguš *et al*. (2024) have argued that coda vowel quality is most likely actively controlled because it takes on one of two discrete values (*a* or *i*) and there is little mixing of different click qualities within a coda. If the number of resonant frequencies (and hence the coda quality) were an automatic artifact not controlled by the whales, we would not expect a correlation between coda type and quality. As such, the uneven distribution of coda qualities across types further supports the claim that coda vowel features are deliberately controlled. The observed interaction between coda type and quality is highly reminiscent of the interaction between tone and segmental vowel features in human languages. In Standard Slovenian, for example, tense vowels, such as *e*, surface preferably with the high tone (produced with more frequent glottal pulses), while lax vowels, such as *ε*, are preferred with the low tone (produced with less frequent glottal pulses).^30,31^

### 4.2 Intrinsic coda vowel duration

Next, we show that there are intrinsic durational differences between the two whale coda vowels. To do so, we analyze focal codas of the 1+1+3 type produced by four whales: Atwood, Fork, Pinchy, and TBB. Since the click count and timing patterns that define the traditional coda types necessarily come with inherent durational differences, the lengths of *a* and *i* intrinsic to those vowel qualities themselves can be compared in a valid manner only within the same coda type. We have chosen the 1+1+3 coda type because this is by far the most common type produced by members of the EC1 clan and the only type that has enough instances of the two coda vowels attested. Additionally, we restrict our analysis to codas produced by four whales only. Atwood, Fork, Pinchy, and TBB each have 50 or more focal codas of the 1+1+3 type captured clearly enough to permit reliable vowel quality annotations. These four included whales have produced 88.5% of all recorded 1+1+3 codas in this dataset. (The other eleven whales have 27 or fewer 1+1+3 codas per whale.) The total number of codas included in this analysis was 628.

The raw distribution of durations for each vowel in Figure 3a,b suggests that the *a*-vowels are intrinsically longer than the *i*-vowels. To confirm this observation, we fit a mixed-effects linear regression model with the duration of the 1+1+3 coda as the dependent variable and the vowel identity as the predictor with a random intercept and slope for the vowel identity for each of the four whales. The *a*-vowels are significantly longer than the *i*-vowels (*β* = − 0.13, *t* = − 6.84).^8^

**Figure 3.**
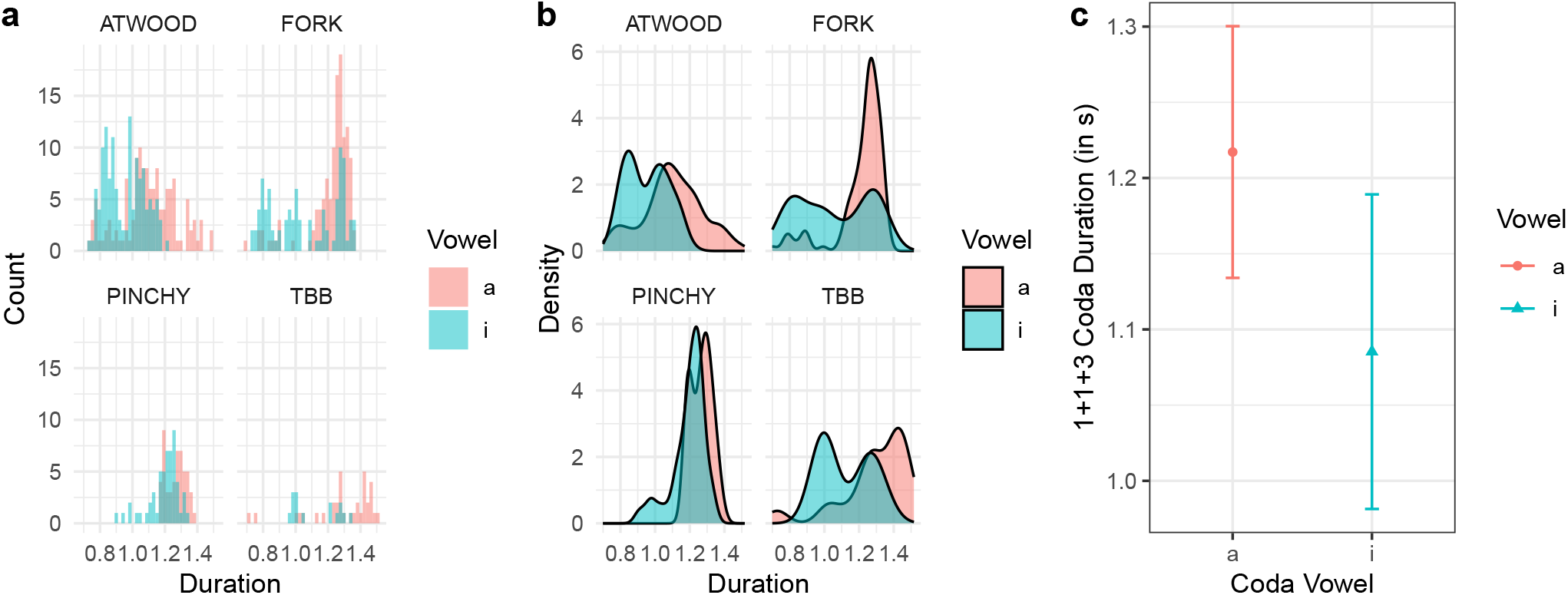
A histogram (**a**) and a density plot (**b**) of raw coda durations (in seconds) of the 1+1+3 coda on four whales. Estimates of the mixed effects linear regression model from Table 2 with 95% CIs (**c**).

This finding has a parallel in human languages, as human vowels have intrinsic durational differences, too. For example, low vowels such as *a* are cross-linguistically longer than high vowels, such as *i* or *u*.^32–34^In the case of human language, the cause of these intrinsic durational differences is articulatory: low vowels, such as *a*, involve a wider opening of the jaw, which takes more time than the restricted movement required for *i* or *u*.^35,36^Since little is known about the precise articulatory mechanics of the whale vocal apparatus, we refrain from speculating about the factors responsible for the intrinsic durations of whale vowels.

**Table 2.** Coefficients of the mixed effects linear regression model with the absolute coda duration (in seconds) as the dependent variable and manually annotated coda vowel type as the dependent variable (treatment-coded with *a* as the base level). The model includes a random intercept for whale identity and a per-whale random slope for coda vowel identity. We take *t*-values above 2.0 as a significant outcome.^7^

|  | estimate | std. error | t-value |
| --- | --- | --- | --- |
| (intercept) | 1.22 | 0.04 | 28.76 |
| vowel *i* | -0.13 | 0.02 | -6.84 |

### 4.3 Contrastive coda vowel length

Based on the raw visualization of coda duration across the two coda vowel qualities (*a* and *i*) in Figure 3, we observe that while the *a*-coda durations are uniformly distributed, the *i*-codas have a bimodal distribution. This suggests that, on top of the intrinsic durational differences between *a* and *i*, the whale communication system differentiates between short *i*-vowels and long *ī*-vowels.

To test this hypothesis, we have run Hartigan’s dip test^37^ for each of the four whales. The test is run separately because different whales have different baseline vowel durations (to be discussed in subsection 4.4). This prevents the potential confound where apparent bimodality stems from whale identity. Table 3 shows that the *a*-vowel duration is unimodal for all four whales, while the *i*-vowel duration for bimodal for at least 3 out of 4 whales. When codas for all four whales are pooled together, the multimodality test remains bimodal for *i*-, but not *a*-coda vowels which parallels the results in individual whales. As in subsection 4.2, the total number of codas included in the analysis was 628. The results suggest that the whale communication system has a contrast between short *i*-codas and long *ī-*codas. A bimodal distribution of *i* vs. *ī* has not been observed for Pinchy. This absence might mean that not all whales produce the length distinction on *i*, or it may be a sampling gap in the recorded dataset. Notably, however, we have no evidence for a bimodal distribution of the duration of *a*.

**Table 3.**
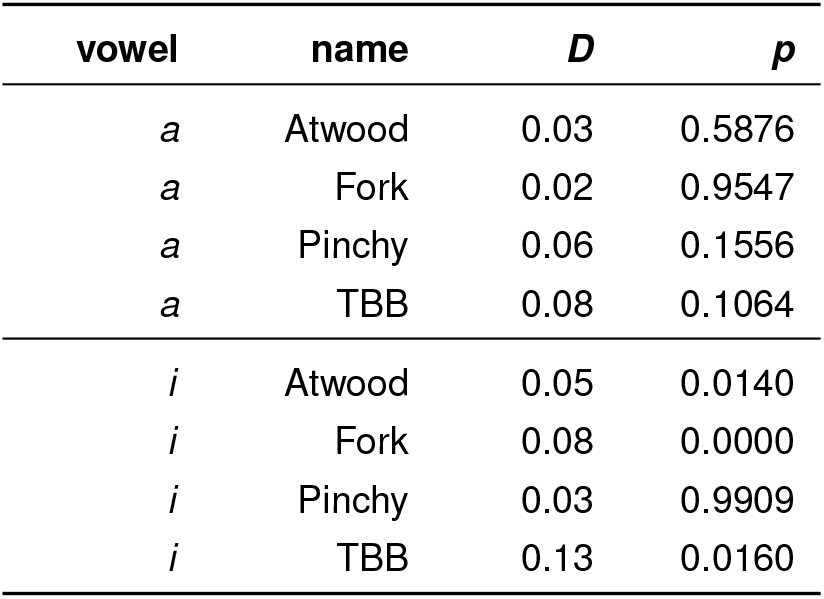
Hartigan’s dip tests for each distribution of coda durations per coda vowel type and each individual whale.^7^.

| vowel | name | *D* | *p* |
| --- | --- | --- | --- |
| *a* | Atwood | 0.03 | 0.5876 |
| *a* | Fork | 0.02 | 0.9547 |
| *a* | Pinchy | 0.06 | 0.1556 |
| *a* | TBB | 0.08 | 0.1064 |
| *i* | Atwood | 0.05 | 0.0140 |
| *i* | Fork | 0.08 | 0.0000 |
| *i* | Pinchy | 0.03 | 0.9909 |
| *i* | TBB | 0.13 | 0.0160 |

It has been known before that codas may have different durations. Gero *et al*. (2016), for example, distinguish between long and short 5R codas.^9^ Sharma *et al*. (2023) categorically cluster coda durations (termed *tempo types*) and propose that codas can have up to five tempo types.^14^ Our contribution is to show that some of the durational distinctions are correlated with the two identified coda vowel qualities (*a* vs. *i*). Unlike previous work, we restrict our analysis to within-type duration, as different coda types have different intrinsic lengths. (This is the case for different tonal contours in human languages as well.^38^) The existence of a pattern where the *i*-coda vowel has two durational subcategories (the short *i* and the long *ī*), whereas the *a*-coda vowel only has one category, means that whales can control both the coda vowel quality as well as coda vowel duration, and that at least a portion of the previously observed length differences are due to different coda vowel qualities.

The distinction between long and short vowels has a clear parallel in human languages, which often use vowel length to distinguish between different words. In Hungarian, for example, the short *bor* means ‘wine,’ while the long *bór* means ‘boron.’ Other languages with contrastive vowel length include Arabic, Finnish, Latin, German, Xhosa, Slovak, and many others.

### 4.4 Baseline coda vowel duration

In subsection 4.2, we demonstrated that the two coda vowels might have intrinsic durational differences. In this section, we show that individual whales also have different baseline vowel durations.

Figure 4 and Table 4 provide random intercepts and slopes for each analyzed whale. We observe that the baseline duration of the *a*-vowel for Atwood is 1.11s, but 1.28s for Pinchy. In other words, the *a*-codas of the 1+1+3 type are on average 170 ms longer for Pinchy than for Atwood. This difference in duration between Pinchy and Atwood is larger than the difference between the intrinsic durations of *a*-codas and *i*-codas. The random slopes also differ among individual whales, but the magnitudes of those differences are smaller. For example, the 1+1+3 *i*-codas are on average 150 ms shorter than the 1+1+3 *a*-codas for Atwood, but only 100 ms shorter for Pinchy. The random slope is negative in all four analyzed whales.

**Table 4.**
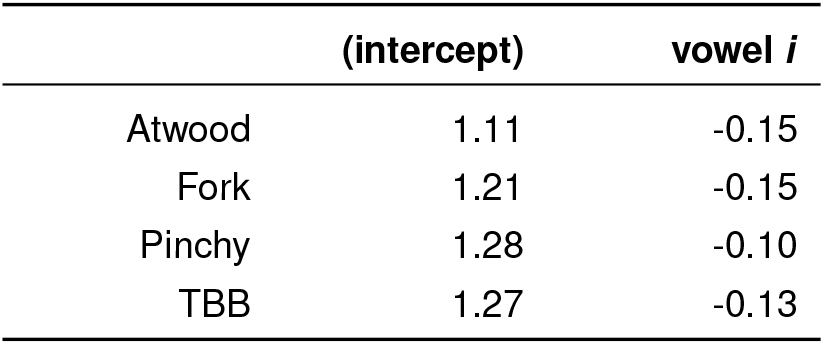
Random intercepts and slopes for the model in Table 2.^7^.

|  | (intercept) | vowel *i* |
| --- | --- | --- |
| Atwood | 1.11 | -0.15 |
| Fork | 1.21 | -0.15 |
| Pinchy | 1.28 | -0.10 |
| TBB | 1.27 | -0.13 |

**Figure 4.**
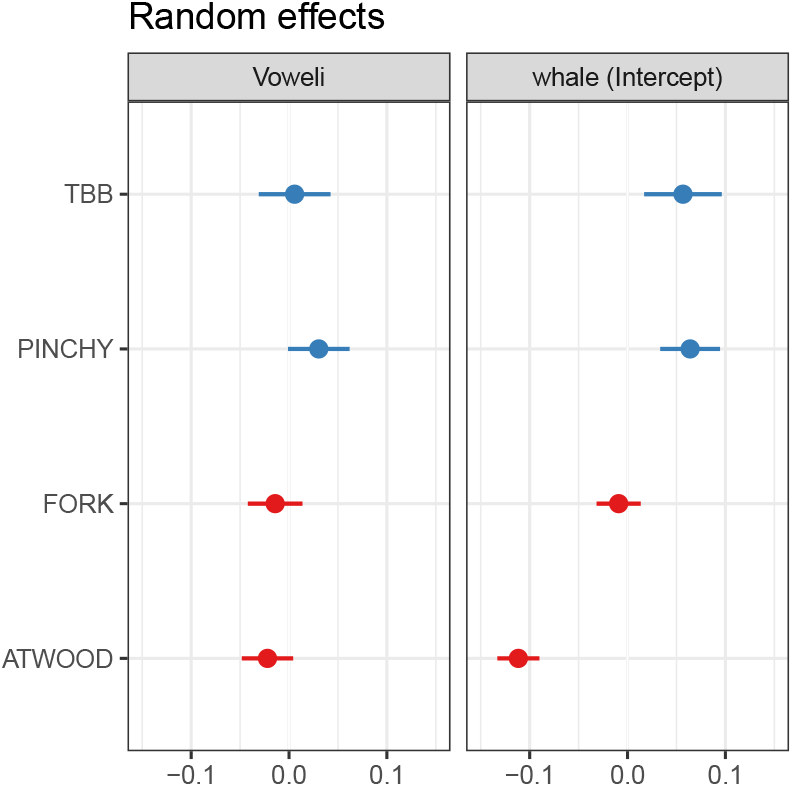
Random slopes (left) and intercepts (right) for each of the four whales for the model in Table 2.]

Sharma *et al*. (2023) identify *rubato*, i. e. tempo modulation across consecutive codas, as one of the parameters that constitute different coda types.^14^ Further, they showed that individuals accurately match coda durations across exchanges. This may have some bearing on driving the baseline differences in duration we observe here. However, in Figure 4, we see that random slopes are rather uniform. If rubato were exclusively responsible for the coda duration differences among individual whales, we would expect random slopes not to be uniform. As such, the baseline click rate we observe is likely not exclusively attributable to the influence of rubato in the sample.

The differences in coda duration among individual whales are paralleled by a similar variation among the speakers of human languages. Different people have different habitual speaking rates,^39,40^which gives rise to differences in average vowel duration. Other phonetic categories, such as vowel quality, differ among humans from speaker to speaker, too.^41^

### 4.5 Coarticulation

In most cases, all clicks within a coda match in vowel quality. This is to say, codas typically consist of all *a*-clicks or all *i*-clicks.^15^ Nevertheless, in some codas, the first click does not match the quality of the rest of the coda. These include cases where the first *a*-click is followed by *i*-clicks (illustrated in Figure 5a), as well as codas whose first *i*-click is followed by *a*-clicks (Figure 5b). Here, we show that the mismatches are more structured than previously thought.

**Figure 5.**
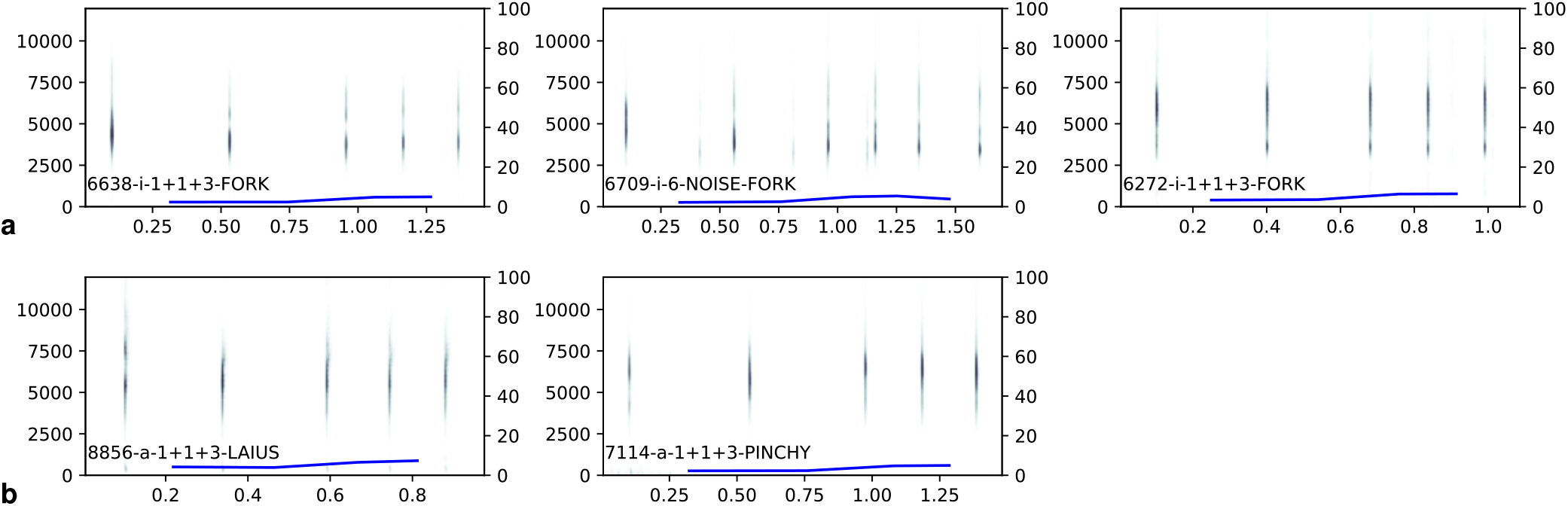
Codas with mismatched first clicks, including “*a*-initial” *i*-codas (**a**), and “*i*-initial” *a*-codas (**b**).^2, 5^

We hypothesize that the quality of the first click is often influenced by the previous coda. As such, we expect to see more mismatches whenever a coda is preceded by a coda of a different quality (i. e. *i* preceded by *a*, or *a* preceded by *i*). To test this hypothesis, we have analyzed mismatches as produced by four whales (Atwood, Fork, Pinchy, and TBB). We included codas of all types (not only 1+1+3). Codas were hand-labeled as “*a*” or “*i*.” Additionally, the first click of each coda was automatically classified as having one or two peaks (i. e. the “*a*” or “*i*” quality) using a simple peak-finding algorithm.^9^ If the coda quality (as hand-labeled) was different from the first click quality (as determined by the algorithm), i. e. if the first click of an *a*-coda was *i* or the first click of an *i*-coda was *a*, we counted that coda as having a “mismatch.” Otherwise, no mismatch was reported. Finally, we limit the analysis to only those cases in which two consecutive codas are at most 10 seconds apart. (If there is a gap of 10 or more seconds between codas, they are annotated as belonging to different *bouts*.) In total, 760 codas were included in this analysis.

We fit the data to a mixed-effects logistic regression model with three predictors: (i) the presence or absence of coda vowel change (if two subsequent focal codas change from *a* to *i* or *i* to *a*), (ii) the quality of the coda vowel with or without the mismatched first click, and (iii) the vowel quality of the preceding coda. The random effects structure includes a random intercept for each of the four whales. We find that the first click is significantly more often mismatched when the whale makes a transition between vowels of two different qualities in sequential codas (from *a* to *i*, or from *i* to *a*), compared to when no change occurs (*β* = 1.31, *z* = 3.07, *p* = 0.002). Mismatched first clicks are also more frequent on *i*-codas than on *a*-codas. The preceding vowel does not have a significant effect on the rate of mismatched first clicks. (I. e., there is no effect of whether the preceding vowel is *i* and the following is *a* or the preceding vowel is *a* and the following is *i*.) Figure 6 illustrates the effects. The coefficients are given in Table 5.

**Table 5.** Estimates of the mixed-effect model in Figure 6.^7^.

|  | estimate | std. error | z-value | Pr(>\|z\|) |
| --- | --- | --- | --- | --- |
| (intercept) | -2.48 | 0.45 | -5.48 | 0.0000 |
| no change | -1.31 | 0.43 | -3.07 | 0.0021 |
| vowel *i* | 1.83 | 0.42 | 4.34 | 0.0000 |
| prev vowel *i* | 0.59 | 0.42 | 1.38 | 0.1667 |

**Figure 6.**
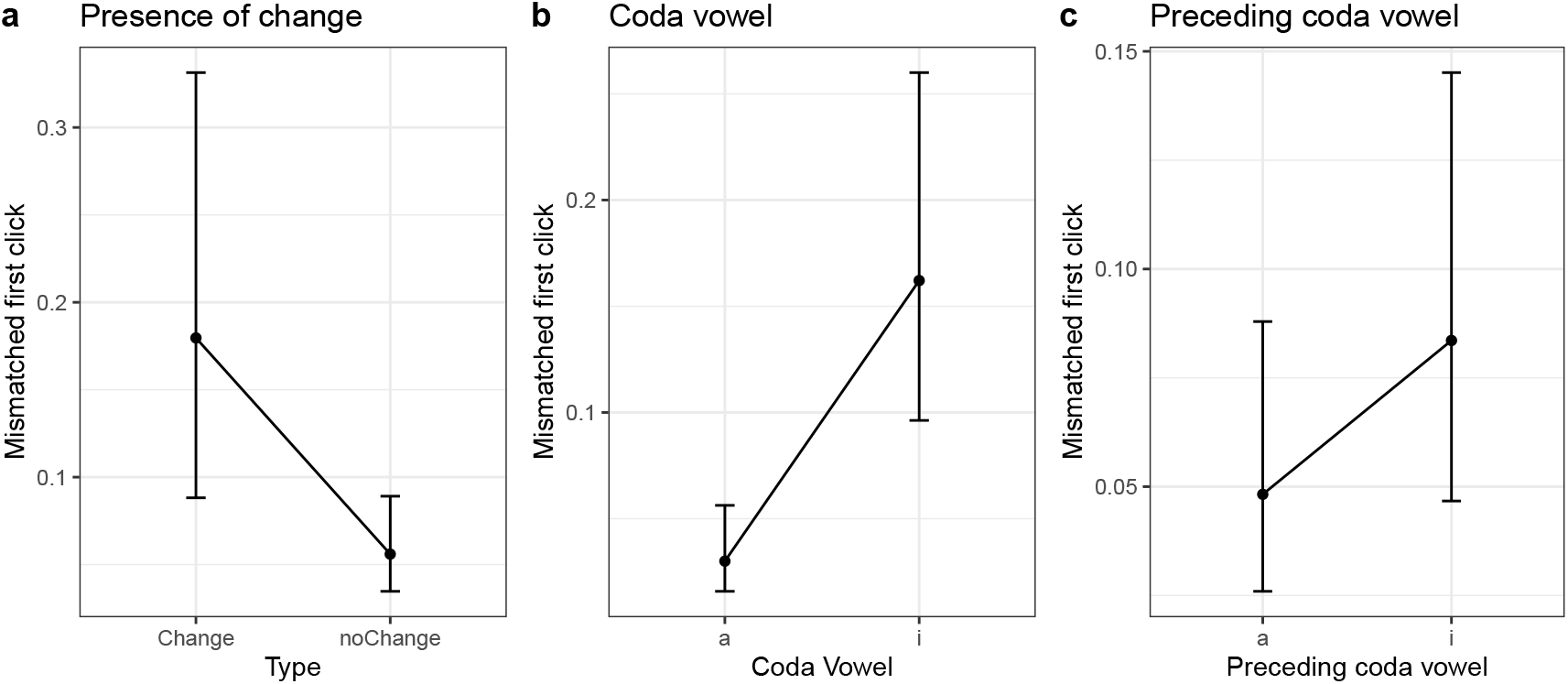
Effects of change, coda vowel, and preceding coda vowel on first-click mismatches. There are significantly more mismatches when the preceding coda has a different vowel quality (**a**). There are significantly more mismatches on *i*-codas than on *a*-codas (**b**). The quality of the preceding vowel does not have a significant effect on the mismatch rate (**c**).

The fact that mismatches are more frequent when changing between different vowel qualities suggests that they result from articulatory constraints on adjusting the acoustic filter responsible for resonant frequencies. This is highly reminiscent of human language coarticulation—a pervasive and versatile phenomenon where the realization of individual sounds is influenced by adjacent segments. For example, the pronunciation of the first vowel in words such as *eli* and *ebi* differs from the pronunciation of the same vowel in *ela* and *eba*. In *eli* and *ebi*, the initial *e* is more close in anticipation of the following *i*, while in *ela* and *eba*, the vowel *e* is realized as more open in the anticipation of *a*.^43–45^Thus, we observe that the whale communication system parallels human languages in the way individual phones affect each other’s articulation.

Finally, we note that the mismatches we observe are mostly limited to the first clicks in a coda. Beguš *et al*. (2024) observe that in some diphthongs, only the first click has a substantially lower or higher frequency. This pattern is clearly seen in Figure 1c.^15^ Both findings suggest that the first click in a coda has a special phonological status.

## 5 Discussion and conclusions

In conclusion, we have investigated five distributional properties of the whale communication system. All five have close parallels in the phonological and phonetic systems of human language. First, we have found that there is a correlation between coda type vowel quality. In the 1+1+3 type, approximately half of the codas are *a*-vowels, while the other half *i*-vowels. In other coda types, such as 5R_1_ or 5R_2_, the *a*-vowels predominate. This has a parallel in some languages, as tones may preferentially co-occur with some vowel qualities.^30^ Second, we have demonstrated that *a*-codas are on average longer than *i*-codas. In human languages, lower vowels are intrinsically longer than short vowels.^35,36^Third, we have shown *i*-codas show a contrast between short (*i*) and long (*ī*) vowels. This closely echoes vowel length distinctions widely attested across the languages of the world. Fourth, our analyses reveal that the whale vocalizations tend to be somewhat faster or slower depending on the whale. Similarly, humans have different habitual speaking rates.^39,40^Finally, we have observed that coda onsets (i. e. first clicks) are sometimes impacted by the quality of the immediately preceding coda. This is an effect of coarticulation, which is also pervasive in human speech.^43–45^

The whale communication system has been previously established to distinguish between coda types that differ in the number of clicks and inter-click intervals.^13^ It has also been shown that codas have structured spectral properties, distinguishing the number of formants and formant trajectories.^15^ Our results indicate that the sperm whale codas not only acoustically resemble, but also behave in ways parallel to human vowels. The sum of all these findings (Table 6) strongly suggests that features of the sperm whale communication system closely parallel many of the phonetic and phonological properties of human language.

**Table 6.** Comparison between features of the whale communication system and human languages.

| sperm whale communication system feature | human language feature | linguistic role | example |
| --- | --- | --- | --- |
| <u>WEILGART, L. & WHITEHEAD, H. (1993),<sup>13</sup> BEGUŠ, G. et al. (2024)<sup>15</sup></u> |  |  |  |
| • coda type (1+1+3, 5R<sub>1</sub>, 5R<sub>2</sub>, etc.) | F0 contour and duration | tonal contour with duration | Vietnamese *sac, nang, nga, hoi, ngang*, vs. *huyen* tones<sup>26</sup> |
| – inter-click intervals | F0 contour | tonal contour | (tonal contrasts) |
| – number of clicks | duration | vowel duration intrinsic to tonal contour | Vietnamese short (*sac, nang*) vs. long (*nga, hoi*) tones<sup>26</sup> |
| <u>BEGUŠ, G. et al. (2024)<sup>15</sup></u> |  |  |  |
| • formant structure | formant structure | vowel quality(-ies) (or nasality) |  |
| – formant number (*i*-coda vs. *a*-coda vowels) | formant number | vowel quality (or nasality) | English *sat* vs. *sit* (or French *beau* vs. *bon*) |
| – formant trajectory | formant trajectory | vocalic target number (mono- vs. diphthongs) | English *sit* vs. *site* |
| <u>PRESENT CONTRIBUTION</u> |  |  |  |
| • correlation coda vowel quality and coda type (§4.1) | correlation between vowel quality and tone |  | no falling tone on lax mid vowels in Slovenian<sup>30</sup> |
| • *a*-codas are on average longer than *i*-codas (§4.2) | vowel duration intrinsic to vowel quality |  | English *sit* vs. *sat* |
| • bimodal distribution of *i*-codas vs. *ī*-codas (§4.3) | bimodal distribution of short and long vowels | phonological vowel length | Hungarian *bor* ‘wine’ vs. *bór* ‘boron’ |
| • correlation between coda length and whale (§4.4) | baseline durational differences across speakers |  | fast-speaking people vs. slow-speaking people |
| • mismatched 1st clicks (§4.5) | coarticulation | integration of consecutive phonological units<sup>43</sup> | *e* in *ebi* higher than in *eba*<sup>44</sup> |

Many previous studies of animal communication systems focused on the analogies with human syntax.^46^ Cotton-top tamarins (*Saguinus oedipus*), for example, can learn local regularities and transitional probabilities between adjacent linguistic elements, but show no evidence of acquiring human-like context-free recursive phrase-structure grammars.^47^ European starlings (*Sturnus vulgaris*), on the other hand, have been argued to recognize recursive center-embedded structures generated by context-free grammars^48^ (although the findings have been hotly contested).^49,50^The songs of humpback whales (*Megaptera novaeangliae*) show structural constraints with periodic characteristics, suggesting a degree of syntax-like organization.^51^

Other animals have also been shown to acquire aspects of their vocalization systems.^46^ This includes other cetaceans, such as toothed whales (*Odontocetes*), e. g. bottlenose dolphins (*Tursiops*), and baleen whales (*Mysticetes*), which can imitate sounds, show “dialectal” variation, or learn new vocalizations over time,^52^ as well as pinnipeds,^53^ with different geographical “variants” reported for—among others—earless seals (*Phocidae*).^54,55^Outside of marine mammals, vocal learning has also been observed in birds. Zebra finches (*Taeniopygia guttata*), for example, have a greater ability to learn songs early in their lives, showing a human-like critical period of “language” acquisition, and—like humans—depend heavily on imitating the adults’ input.^56^

Little attention has been devoted to the investigation of phonological abilities in non-human animals.^46^ Some studies focused on whether animals can attune to the features of human phonologies. E. g., tamarins have been shown to discriminate between sentences in Dutch (a stress-timed language) and Japanese (a syllable-timed language), which suggests they can recognize human rhythmic categories.^57^ European starlings (*Sturnus vulgaris*) can learn the prototypical realization of the vowels in *heat* and *hit*, showing a “perceptual magnet effect” responsible for the organization of human language categories.^58^ Fewer studies have been conducted on the phonological properties inherent in actual animal communication systems.^46^ Some similarities to human phonology have been observed, but the findings are limited. Bird song involves the modulation of fundamental frequency (F0),^56^ similarly to human tonal contours and whale coda types. Bird songs and humpback whale songs both show hierarchical organization, which has often been compared to human languages. For example, individual bird song “notes” have been analogized to *phonemes*, and motifs (note clusters) to higher-level prosodic constituents, such as *syllables* and *phrases*.^51,56,59–61^The songs of northern mockingbirds (*Mimus polyglottus*) and humpback whales include structured repetitions, which pattern in ways that resemble human rhyming.^62–64^African elephant (*Loxodonta africana*) rumbles can be produced via an oral or a nasal route, giving rise to two different discrete formant structures,^65^ in a way analogous to the distinction between human oral (*a, b*) and nasal (*ã, m*) phones.

Against this background, we show that the sperm whale communication system has previously undocumented characteristics that make it similar to human phonology. Sperm whale vocalizations make use of a genuine source-filter mechanism, where phonic lip clicks are responsible for the fundamental frequency of codas, and the distal air sac—for the modulation of resonant frequencies.^15^ This is analogous to the way that the frequency of human vocal fold vibrations gives rise to pitch, and the vocal tract position alters vowel quality. On top of that, the sperm whale codas instantiate one of two discrete vowel qualities: each coda is either an *a*-coda or *i*-coda, with no in-between values. Human phoneme inventories are organized in a similarly discrete fashion: Even though the resonant frequencies (F1 and F2) can be modulated continuously, each language makes use only of a finite set of vowels. A subset of the *a*-coda vowels shows a formant transition, with formant ascending or descending in the course of the coda, in a way analogous to human diphthongs. In the whale communication system, the features of coda type and quality are partially (though not completely) independent, as are tones and vowel qualities in human languages. To the best of our knowledge, no other animal communication system has previously been reported to exhibit structured variability by combining features of fundamental frequency (F0) and formant structure (F1, F2). Taken together, our findings demonstrate that sperm whale vocalizations are highly complex and likely constitute one of the most phonologically sophisticated communication systems in the animal kingdom.

## Data Availability

The spectral data and the code for the analysis and figures are available at http://doi.org/10.17605/OSF.IO/A32KE.

## Materials & Correspondence

Correspondence and material requests should be addressed to G.B.

## Acknowledgements

This study was funded by Project CETI via grants from Dalio Philanthropies and Ocean X; Sea Grape Foundation; and Rosamund Zander/Hansjorg Wyss through The Audacious Project: A collaborative funding initiative housed at TED.

Fieldwork for the Dominica Sperm Whale Project during tagging years in 2014-2018 was supported by an FNU fellow-ship for the Danish Council for Independent Research supplemented by a Sapere Aude Research Talent Award, a Carlsberg Foundation expedition grant, a grant from Focused on Nature, two Explorer Grants from the National Geographic Society, and supplementary grants from the Arizona Center for Nature Conservation, Quarters For Conservation, the Dansk Akustisks Selskab, Oticon Foundation, and the Dansk Tennis Fond all to SG. Further funding was provided by a Discovery and Equipment grant from the Natural Sciences and Engineering Research Council of Canada (NSERC) to Hal Whitehead of Dalhousie University, an FNU large frame grant, and a Villum Foundation Grant to Peter Madsen of Aarhus University.

We are grateful to Kristian Beedholm for CodaSorter, as well as Mark Johnson and Peter L. Tyack for their in-kind contribution of DTAGs and associated code during the research. We thank the Chief Fisheries Officers and the Dominica Fisheries Division officers for research permits and their collaboration in data collection; all the crews of R/V Balaena and the DSWP team for data collection, curation, and annotation; as well as Dive Dominica, Al Dive, and W.E.T. Dominica for logistical support while in Dominica.

## Footnotes

1 Different human words refer to different things, and changing a sound within a word may change its meanings. The study of the whale communication system has not yet revealed the meanings encoded by particular codas or their sequences. We leave the question of the information conveyed by click sequences aside, and draw our comparisons based solely on the *structural* properties of the codas-based communication system.

2 Spectrograms were calculated using *Praat*’s^19^ *To Spectrogram*…function via *Parselmouth*,^20^ with a 5 ms analysis window and maximum frequency of 12000 Hz. Spectrogram values are plotted with *Matplotlib*’s^21^ *imshow* function, using log normalization and masking values below a threshold in order to enhance visibility of the peaks. This threshold was set separately for each spectrogram based on its minimum value plus the standard deviation of its values raised to a constant power.

3 In addition to codas used for communication, sperm whales also use clicks for echolocation. Coda clicks and echolocation clicks are acoustically distinguishable.^22–24^This paper focuses only on the coda clicks used for communication.

4 However, our analogy has a limit: While in human languages, different tones can be associated with different meanings, the meanings conveyed by sperm whale codas have not been established.

5 The pitch plots were calculated as the inverse of each ICI (in seconds) in order to express the period of each click interval in Hz. Plot points are centered within each interval on the time axis.

6 In subsection 4.5, we also use a simple peak-finding algorithm to automatically classify the first click of each coda as having one or two peaks, i. e. the “*a*” or “*i*” quality.

7 The LATEX tables were generated in *R 4*.*3*.*0*^28^ with the *xtable 1*.*8-4*^29^ package. Their typesetting was adjusted manually.

8 Testing this difference on all whales is more challenging because of the sparsity of attested codas on other whales. This makes estimates with random slopes for vowel identity challenging. The quality of data is also lower for whales with fewer codas attested. In some models with per-whale random slopes for vowel identity make the estimate of the vowel identity non-significant. However, when all whales and codas are included in the analysis, the vowel remains a significant predictor for the 1+1+3 coda type when only codas for which both the automatic analysis (whereby all clicks in a coda are of one type) and the hand-annotated analyses agree.

9 The spectrum for each click was calculated by extracting a 3.5 ms audio segment from approximately −2.0 ms preceding the click peak to +1.5 ms after and applying *scipy*’s^42^ *welch* function to estimate spectral power. The *welch* parameters *rate, nperseg*, and *nfft* were respectively set to the audio sample rate (120000 samples/sec), the number of samples in the extracted click (420), and 512. We used *find_peaks* from *scipy* to locate peaks in the spectrum between 1000 and 10000 Hz. The *distance* parameter required the minimum distance between adjacent peaks to be at least 1500 Hz, and *height* ensured each peak was at least 25% of the height of the highest peak in the range. The ordered frequencies of the two highest spectral peaks were recorded as the first and second spectral peaks. The click vowel qualities “*a*” and “*i*” were assigned to each click based on the number of spectral peaks. If there were two or more peaks, the *i*-quality was assigned. If there were fewer peaks, the *a*-quality was assigned. The same algorithm was used in Beguš *et al*. (2024).^15^

## References

1. Whitehead, H. Sperm Whales: Social Evolution in the Ocean (University of Chicago Press, Chicago, 2003).

2. Rendell, L. E. & Whitehead, H. Vocal clans in sperm whales (Physeter macrocephalus). Proc. Royal Soc. London. Ser. B: Biol. Sci. 270, 225–231, DOI: 10.1098/rspb.2002.2239 (2003). https://royalsocietypublishing.org/doi/pdf/10.1098/rspb.2002.2239.

3. Worthington, L. V. & Schevill, W. E. Underwater sounds heard from sperm whales. Nature 180, 291–291, DOI: 10.1038/180291a0 (1957).

4. Watkins, W. A. & Schevill, W. E. Sperm whale codas. The J. Acoust. Soc. Am. 62, 1485–1490, DOI: 10.1121/1.381678 (1977). https://pubs.aip.org/asa/jasa/article-pdf/62/6/1485/11470326/1485_1_online.pdf.

5. Whitehead, H. & Weilgart, L. Patterns of visually observable behaviour and vocalizations in groups of female sperm whales. Behaviour 118, 275 – 296, DOI: 10.1163/156853991×00328 (1991).

6. Andreas, J. et al. Toward understanding the communication in sperm whales. iScience 25, 104393, DOI: 10.1016/j.isci.2022.104393 (2022).

7. Amano, M., Kourogi, A., Aoki, K., Yoshioka, M. & Mori, K. Differences in sperm whale codas between two waters off Japan: Possible geographic separation of vocal clans. J. Mammal. 95, 169–175, DOI: 10.1644/13-MAMM-A-172 (2014). https://academic.oup.com/jmammal/article-pdf/95/1/169/2729464/95-1-169.pdf.

8. Amorim, T. O. S. et al. Coda repertoire and vocal clans of sperm whales in the western atlantic ocean. Deep. Sea Res. Part I: Oceanogr. Res. Pap. 160, 103254, DOI: 10.1016/j.dsr.2020.103254 (2020).

9. Gero, S., Whitehead, H. & Rendell, L. E. Individual, unit and vocal clan level identity cues in sperm whale codas. Royal Soc. Open Sci. 3, 150372, DOI: 10.1098/rsos.150372 (2016). https://royalsocietypublishing.org/doi/pdf/10.1098/rsos.150372.

10. Huijser, L. A. E. et al. Vocal repertoires and insights into social structure of sperm whales (Physeter macrocephalus) in Mauritius, southwestern Indian Ocean. Mar. Mammal Sci. 36, 638–657, DOI: 10.1111/mms.12673 (2020). https://onlinelibrary.wiley.com/doi/pdf/10.1111/mms.12673.

11. Hersh, T. A. et al. Evidence from sperm whale clans of symbolic marking in non-human cultures. Proc. Natl. Acad. Sci. 119, e2201692119 (2022).

12. Rendell, L. E., Mesnick, S. L., Dalebout, M. L., Burtenshaw, J. & Whitehead, H. Can genetic differences explain vocal dialect variation in sperm whales, Physeter macrocephalus? Behav. Genet. 42, 332–343, DOI: 10.1007/s10519-011-9513-y (2012).

13. Weilgart, L. & Whitehead, H. Coda communication by sperm whales (Physeter macrocephalus) off the Galápagos Islands. Can. J. Zool. 71, 744–752, DOI: 10.1139/z93-098 (1993). 10.1139/z93-098.

14. Sharma, P. et al. Contextual and combinatorial structure in sperm whale vocalisations. Nat. Commun. 15, 3617, DOI: 10.1038/s41467-024-47221-8 (2024).

15. Beguš, G., Sprouse, R. L., Leban, A., Silva, M. & Gero, S. Vowels and diphthongs in sperm whales. Manuscript DOI: 10.31219/osf.io/285cs (2024).

16. Gero, S. et al. Behavior and social structure of the sperm whales of Dominica, West Indies. Mar. Mammal Sci. 30, 905–922 (2014).

17. Best, P. B. Social organization in sperm whales, physeter macrocephalus. In Behavior of marine animals: Current perspectives in research, 227–289 (Springer, Boston, USA, 1979).

18. Johnson, M. & Tyack, P. L. A digital acoustic recording tag for measuring the response of wild marine mammals to sound. IEEE J. Ocean. Eng. 28, 3–12 (2003).

19. Boersma, P. & Weenink, D. Praat. Doing phonetics by computer. Version 6.4.28. Software, http://www.praat.org/ (2025).

20. Jadoul, Y., Thompson, B. & de Boer, B. Introducing Parselmouth: A Python interface to Praat. J. Phonetics 71, 1–15, DOI: 10.1016/j.wocn.2018.07.001 (2018).

21. Hunter, J. D. Matplotlib: A 2d graphics environment. Comput. Sci. & Eng. 9, 90–95, DOI: 10.1109/MCSE.2007.55 (2007).

22. Madsen, P. T. et al. Sperm whale sound production studied with ultrasound time/depth-recording tags. J. Exp. Biol. 205, 1899–1906, DOI: 10.1242/jeb.205.13.1899 (2002). https://journals.biologists.com/jeb/article-pdf/205/13/1899/1240975/1899.pdf.

23. Madsen, P., Wahlberg, M. & Møhl, B. Male sperm whale (Physeter macrocephalus) acoustics in a high-latitude habitat: Implications for echolocation and communication. Behav. Ecol. Sociobiol. 53, 31–41, DOI: 10.1007/s00265-002-0548-1 (2002).

24. Møhl, B., Wahlberg, M., Madsen, P. T., Heerfordt, A. & Lund, A. The monopulsed nature of sperm whale clicks. The J. Acoust. Soc. Am. 114, 1143–1154, DOI: 10.1121/1.1586258 (2003). https://pubs.aip.org/asa/jasa/article-pdf/114/2/1143/8092391/1143_1_online.pdf.

25. Duanmu, S. The Phonology of Standard Chinese. The Phonology of the World’s Languages (Oxford University Press, Oxford, 2007), 2nd ed. edn.

26. Alves, M. Tonal features and the development of Vietnamese tones. Work. Pap. Linguist. 27, 1–13 (1995).

27. Fant, G. Acoustic Theory of Speech Production (Mouton, The Hague, 1960).

28. R Core Team. R: A Language and Environment for Statistical Computing. R Foundation for Statistical Computing, Vienna, Austria (2018).

29. Dahl, D. B., Scott, D., Roosen, C., Magnusson, A. & Swinton, J. xtable: Export Tables to LaTeX or HTML (2019). R package version 1. 8–4.

30. Becker, M. et al. Interactions of tone and ATR in Slovenian. In Segmental Structure and Tone, 11–26 (Walter de Gruyter Berlin, 2017).

31. Begus, G. & Jurgec, P. Tone, stress, quantity, and quality: prosodic patterns and tonal wug-tests in Žiri slovenian. PsyArXiv DOI: 10.31234/osf.io/b7uj5 (2025).

32. Heffner, R. S. Notes on the length of vowels. Am. Speech 128–134 (1937).

33. House, A. S. & Fairbanks, G. The influence of consonant environment upon the secondary acoustical characteristics of vowels. The J. Acoust. Soc. Am. 25, 105–113 (1953).

34. Maack, A. Die spezifische Lautdauer deutscher Sonanten. STUF-Language Typology Universals 3, 190–231 (1949).

35. Lehiste, I. Suprasegmentals (MIT, Cambridge, MA, 1970).

36. Fischer-Jørgensen, E. Sound duration and place of articulation. STUF-Language Typology Universals 17, 175–208, DOI: 10.1524/stuf.1964.17.16.175 (1964).

37. Maechler, M. diptest: Hartigan’s Dip Test Statistic for Unimodality - Corrected (2024). R package version 0. 77-1.

38. Nguyêñ, V. L. & Edmondson, J. Tones and voice quality in modern northern Vietnamese: Instrumental case studies. Mon-Khmer Stud. 28, 1–18 (1997).

39. Tsao, Y.-C. & Weismer, G. Interspeaker variation in habitual speaking rate: Evidence for a neuromuscular component. J. Speech, Lang. Hear. Res. 40, 858–866 (1997).

40. Tanner, J. et al. Exploring the anatomy of articulation rate in spontaneous English speech: Relationships between utterance length effects and social factors. arXiv preprint arXiv:2408.06732 (2024).

41. Johnson, K. A. Processes of speaker normalization in vowel perception (The Ohio State University, 1988).

42. Virtanen, P. et al. SciPy 1.0: Fundamental algorithms for scientific computing in Python. Nat. Methods 17, 261–272, DOI: 10.1038/s41592-019-0686-2 (2020).

43. Kühnert, B. & Nolan, F. The origin of coarticulation. In Hardcastle, W. & Hewlett, N. (eds.) Coarticulation: Theoretical and Empirical Perspectives, 7–30 (CUP, Cambridge, 1999).

44. Scripture, E. W. The Elements of Experimental Phonetics (Charles Scribner’s Sons, New York, 1904).

45. Hardcastle, W. Experimental studies in lingual coarticulation. In Asher, R. & Henderson, E. (eds.) Towards a History of Phonetics, 50–66 (Edinburgh University Press, Edinburgh, 1981).

46. Yip, M. J. The search for phonology in other species. Trends Cogn. Sci. 10, 442–446, DOI: 10.1016/j.tics.2006.08.001 (2006).

47. Fitch, W. T. & Hauser, M. D. Computational constraints on syntactic processing in a nonhuman primate. Science 303, 377–380, DOI: 10.1126/science.1089401 (2004). https://www.science.org/doi/pdf/10.1126/science.1089401.

48. Gentner, T. Q., Fenn, K. M., Margoliash, D. & Nusbaum, H. C. Recursive syntactic pattern learning by songbirds. Nature 440, 1204–1207 (2006).

49. Corballis, M. C. Recursion, language, and starlings. Cogn. Sci. 31, 697–704 (2007).

50. Beecher, M. D. Why are no animal communication systems simple languages? Front. Psychol. 12, DOI: 10.3389/fpsyg.2021.602635 (2021).

51. Suzuki, R., Buck, J. R. & Tyack, P. L. Information entropy of humpback whale songs. The J. Acoust. Soc. Am. 119, 1849–1866, DOI: 10.1121/1.2161827 (2006). https://pubs.aip.org/asa/jasa/article-pdf/119/3/1849/14876116/1849_1_online.pdf.

52. Janik, V. & Slater, P. J. B. Vocal learning in mammals. Adv. Study Behav. 26, 59–99, DOI: 10.1016/s0065-3454(08)60377-0 (1997).

53. Fitch, W. T. The biology and evolution of music: A comparative perspective. Cognition 100, 173–215, DOI: 10.1016/j.cognition.2005.11.009 (2006).

54. Cleator, H. J., Stirling, I. & Smith, T. G. Underwater vocalizations of the bearded seal (Erignathus barbatus). Can. J. Zool. 67, 1900–1910, DOI: 10.1139/z89-272 (1989). 10.1139/z89-272.

55. Thomas, J. A. & Golladay, C. L. Geographic variation in leopard seal (Hydrurga leptonyx) underwater vocalizations. In Kastelein, R., Thomas, J. A. & Nachtigall, P. E. (eds.) Sensory Systems of Aquatic Mammals, 201–221 (De Spil Publishers, Woerden, The Netherlands, 1996).

56. Doupe, A. J. & Kuhl, P. K. Birdsong and human speech: Common themes and mechanisms. Annu. Rev. Neurosci. 22, 567–631, DOI: 10.1146/annurev.neuro.22.1.567 (1999).

57. Ramus, F., Hauser, M. D., Miller, C., Morris, D. & Mehler, J. Language discrimination by human newborns and by cotton-top tamarin monkeys. Science 288, 349–351, DOI: 10.1126/science.288.5464.349 (2000). https://www.science.org/doi/pdf/10.1126/science.288.5464.349.

58. Kluender, K. R., Lotto, A. J., Holt, L. L. & Bloedel, S. L. Role of experience for language-specific functional mappings of vowel sounds. The J. Acoust. Soc. Am. 104, 3568–3582, DOI: 10.1121/1.423939 (1998). https://pubs.aip.org/asa/jasa/article-pdf/104/6/3568/10679219/3568_1_online.pdf.

59. Gentner, T. Q. & Hulse, S. H. Perceptual mechanisms for individual vocal recognition in european starlings, Sturnus vulgaris. Animal behaviour 56, 579–594, DOI: 10.1006/anbe.1998.0810 (1998).

60. Payne, R. S. & McVay, S. Songs of humpback whales. Science 173, 585–597, DOI: 10.1126/science.173.3997.585 (1971). https://www.science.org/doi/pdf/10.1126/science.173.3997.585.

61. Gentner, T. Q. & Hulse, S. H. Perceptual classification based on the component structure of song in European starlings. The J. Acoust. Soc. Am. 107, 3369–3381, DOI: 10.1121/1.429408 (2000). https://pubs.aip.org/asa/jasa/article-pdf/107/6/3369/12240664/3369_1_online.pdf.

62. Payne, K. The progressively changing songs of humpback whales: A window on the creative process in a wild animal. In Freeman, W., Wallin, N. L., Merker, B. & Brown, S. (eds.) The Origins of Music, 135–150, DOI: 10.7551/mitpress/5190.001.0001 (The MIT Press, Cambridge, MA, 2000).

63. Guinee, L. N. & Payne, K. B. Rhyme-like repetitions in songs of humpback whales. Ethology 79, 295–306, DOI: 10.1111/j.1439-0310.1988.tb00718.x (1988). https://onlinelibrary.wiley.com/doi/pdf/10.1111/j.1439-0310.1988.tb00718.x.

64. Thompson, N. S. et al. Variation in the bout structure of northern mockingbird (Mimus polyglottus) singing. Bird Behav. 13, 93–98 (2000).

65. Stoeger, A. S. et al. Visualizing sound emission of elephant vocalizations: Evidence for two rumble production types. PLOS ONE 7, 1–8, DOI: 10.1371/journal.pone.0048907 (2012).

